# Human *trans*-editing enzyme displays tRNA acceptor stem specificity and relaxed amino acid selectivity

**DOI:** 10.1101/2020.09.11.293571

**Authors:** Oscar Vargas-Rodriguez, Marina Bakhtina, Daniel McGowan, Jawad Abid, Yuki Goto, Hiroaki Suga, Karin Musier-Forsyth

## Abstract

Accurate translation of genetic information into proteins is vital for cell sustainability. ProXp-ala prevents proteome-wide Pro-to-Ala mutations by hydrolyzing misacylated Ala-tRNA^Pro^, which is synthesized by prolyl-tRNA synthetase (ProRS). Bacterial ProXp-ala was previously shown to combine a size-based exclusion mechanism with conformational and chemical selection for the recognition of the alanyl moiety, while tRNA^Pro^ is selected via recognition of tRNA acceptor stem elements G72 and A73. The identity of these critical bases changed during evolution with eukaryotic cytosolic tRNA^Pro^ possessing a cytosine at the corresponding positions. The mechanism by which eukaryotic ProXp-ala adapted to these changes remains unknown. In this work, recognition of the aminoacyl moiety and tRNA acceptor stem by human (*Hs*) ProXp-ala was examined. Enzymatic assays revealed that *Hs* ProXp-ala requires C72 and C73 in the context of *Hs* cytosolic tRNA^Pro^ for efficient deacylation of mischarged Ala-tRNA^Pro^. The strong dependence on these bases prevents cross-species deacylation of bacterial Ala-tRNA^Pro^ or of *Hs* mitochondrial Ala-tRNA^Pro^ by the human enzyme. Similar to the bacterial enzyme, *Hs* ProXp-ala showed strong tRNA acceptor-stem recognition but differed in its amino acid specificity profile relative to bacterial ProXp-ala. Changes at conserved residues in both the *Hs* and bacterial ProXp-ala substrate binding pockets modulated this specificity. These results illustrate how the mechanism of substrate selection diverged during the evolution of the ProXp-ala family and provides the first example of a *trans*-editing domain whose specificity evolved to adapt to changes in its tRNA substrate.

## Introduction

Correct pairing of amino acids with their corresponding tRNAs by aminoacyl-tRNA synthetases (ARSs) is fundamental for faithful translation of genetic information into proteins. These enzymes are divided into two classes based on structural differences within their aminoacylation active sites. The chemical and structural similarities of certain classes of amino acids challenge the ability of the synthetic active site, wherein aminoacyl-adenylate formation occurs, to properly select only cognate amino acids at this first step. Consequently, ARSs frequently ligate tRNAs with the wrong amino acid during the second step of aminoacylation (1). If left uncorrected, misacylation may lead to the incorporation of amino acids in response to the wrong codon (i.e., mistranslation); accumulation of these errors has been linked to cell defects in diverse organisms (1-5). To prevent mistranslation, many ARSs have acquired quality control mechanisms that enable deacylation of tRNAs immediately after their synthesis in a reaction known as post-transfer editing. This reaction may occur in *cis* in a domain distinct from the aminoacylation active site and seven members of both classes of ARSs possess such an editing domain (1). Class II synthetases have additionally been shown to edit aminoacyl-tRNAs in *trans*, that is, following substrate dissociation and rebinding (6-8). In addition, free-standing editing domains that are evolutionally related to the editing domains of three class II ARSs have been identified in all Domains of life; these enzymes lack aminoacylation activity and catalyze deacylation exclusively in *trans* (8).

The critical role of aminoacyl-tRNA editing is highlighted by the different phenotypes associated with editing deficiencies (9-11). For example, murine cells harboring an editing-deficient transgene for valyl-tRNA synthetase transition to an apoptotic-like state due to the incorporation of Thr at Val codons (2), while a mild defect in the editing activity of AlaRS provokes ataxia in mice (3). In *E. coli*, severe oxidative conditions inactivate the editing activity of threonyl-tRNA synthetase, which impairs cell growth (12), whereas heritable genome mutations have been identified in aging *E. coli* cells resulting from accumulation of translational errors from an editing defective isoleucyl-tRNA synthetase (13).

Prolyl-tRNA synthetases (ProRSs) commonly misactivate Ala and Cys, which are smaller or similar in size to cognate Pro (14-17). In most bacteria, mistranslation of Pro codons with Ala is prevented by the ProRS editing domain, which is known as the insertion (INS) domain due to its location between the class II consensus Motifs 2 and 3 (18). This domain specifically hydrolyzes Ala-tRNA^Pro^ but rejects Pro- and Cys-tRNA^Pro^ (14). The later error is corrected by a single-domain *trans*-editing enzyme known as YbaK, which is homologous to the INS domain but possesses unique specificity for Cys-tRNAs (19). While tRNA^Pro^ misacylation with Ala is an inherent characteristic of ProRSs, the INS domain is not present in some bacteria nor is it found in most eukaryotes (15). Instead, many organisms that lack INS encode ProXp-ala, another homolog of the INS domain that hydrolyzes Ala-tRNA^Pro^ in *trans* (20-22). Structural studies revealed that the INS domain from *Enterococcus faecium* and ProXp-ala from *Caulobacter crescentus* (*Cc*) share a conserved tertiary structure and key catalytic residues (23). Biochemical studies revealed that *Cc* ProXp-Ala and *Escherichia coli* (*Ec*) ProRS share similar size-exclusion and chemical-based mechanisms for recognition of the aminoacyl moiety, limiting their ability to hydrolyze tRNA^Pro^ charged with cognate Pro as well as other amino acids that are larger than Ala (24,25). In contrast to their shared mechanism of amino acid recognition, INS and bacterial ProXp-ala developed distinct strategies for tRNA^Pro^ selection. The INS domain lacks intrinsic tRNA specificity and relies on the recognition of the tRNA anticodon by the anticodon binding domain of ProRS, whereas *Cc* ProXp-ala displays strong recognition of the acceptor stem elements of bacterial tRNA^Pro^ (26). The tRNA specificity of both enzymes is crucial for preventing deacylation of cognate Ala-tRNA^Ala^.

Bacterial tRNA^Pro^ sequences show high conservation of the A73 discriminator base and the C1:G72 base pair, which is unique to the tRNA^Pro^ acceptor stem (27). *Cc* ProXp-ala recognizes tRNA^Pro^ through specific interactions with G72 and A73 (26). These bases are also critical for aminoacylation of tRNA^Pro^ by bacterial ProRS (28); thus, they play a dual role in ensuring the correct aminoacylation of tRNA^Pro^ with Pro and in editing of misacylated Ala-tRNA^Pro^. The identity of these bases at the top of the tRNA^Pro^ acceptor stem changed during evolution; eukaryotic cytosolic (cyto) tRNA^Pro^ encodes C73 and G1:C72 (Fig. 1A) (27). These differences in the acceptor stem prevent aminoacylation of human (*Hs*) cyto tRNA^Pro^ by bacterial ProRS (29) as well as deacylation of *Hs* cyto Ala-tRNA^Pro^ (hereafter referred to as *Hs* Ala-tRNA^Pro^) by bacterial ProXp-ala (26). Even though bacterial tRNA^Pro^ is a poor substrate for *Hs* ProRS, the human enzyme only weakly relies on recognition of the tRNA^Pro^ acceptor-stem bases, while strongly relying on interactions with the anticodon, as well as other structural features of *Hs* tRNA^Pro^ (30). A previous study using a variant of tRNA^Pro^ that could readily be misacylated with Ala by alanyl-tRNA synthetase (AlaRS) concluded that *Hs* ProXp-ala has amino acid specificity but lacks tRNA specificity (31). Here, we prepared native aminoacyl-tRNA^Pro^ substrates to investigate the amino acid and tRNA specificity of *Hs* ProXp-ala. Our findings suggest that the substrate binding pocket of *Hs* ProXp-ala displays different amino acid selectivity relative to that of the bacterial enzyme, but the human enzyme maintains strong acceptor stem recognition. We also identified residues in the substrate binding pocket of both bacterial and *Hs* ProXp-ala that modulate their amino acid specificity. The results reported here provide insights into the evolutionary relationship between class II aminoacyl-tRNA synthetases and editing domains.

**Figure 1.**
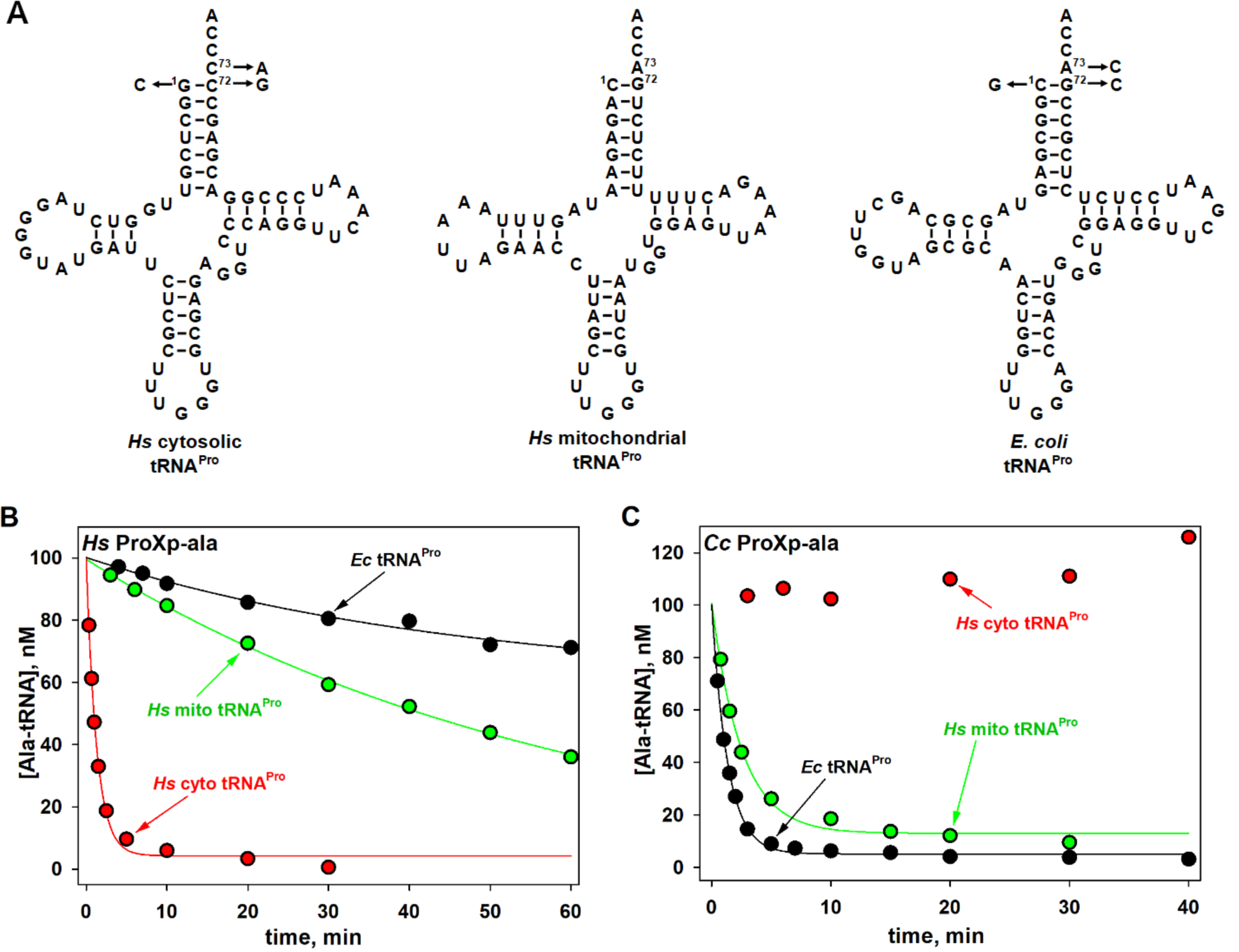
Cross-species deacylation of *Hs* and *Cc* ProXp-ala. (A) Predicted secondary structures of *Hs* cytosolic (cyto), mitochondrial (mito) and *Ec* tRNA^Pro^. Arrows indicate the acceptor stem positions mutated in this study. (B) Deacylation of *Hs* cyto, mito, and *Ec* Ala-tRNA^Pro^ by *Hs* ProXp-ala. (C) Deacylation of *Hs* cyto, mito, and *Ec* Ala-tRNA^Pro^ by *Cc* ProXp-ala. Representative time courses of 3 independent experiments are shown. Reactions were performed with 1.5 µM *Hs* ProXp-ala and 100 nM Ala-tRNA^Pro^ at 30 °C (panel B) or with 0.75 µM *Cc* ProXp-ala and 100 nM Ala-tRNA^Pro^ at 18 °C (panel C), as described in the Experimental Procedures.

## Results

### Hs ProXp-ala robustly deacylates Ala-tRNA^Pro^

In this work, we used *in vitro* transcribed *Hs* cyto tRNA^Pro^ (Fig. 1A) and full-length recombinant *Hs* ProXp-ala expressed and purified in *E. coli*. To obtain sufficient yield of misacylated *Hs* Ala-tRNA^Pro^ substrates without the need for tRNA acceptor stem mutation, we used flexizyme technology, which allows tRNAs to be charged with virtually any amino acid (proteinogenic or non-proteinogenic) (Fig. S1) (32). Our initial experiments showed that *Hs* ProXp-ala robustly deacylated Ala-tRNA^Pro^ *in vitro*. Based on an analysis of *Hs* Ala-tRNA^Pro^ deacylation by varying concentrations of *Hs* ProXp-ala under single-turnover conditions, we determined an apparent dissociation constant (K_d_) of 10.2 µM and a maximum rate constant (k_max_) of 3.8 min^-1^ (Fig. 2). Relative to the cyto tRNA, much weaker deacylation of *Hs* mitochondrial (mito) Ala-tRNA^Pro^ by *Hs* ProXp-ala was observed (Fig. 1B). Conversely, *Cc* ProXp-Ala deacylates mito Ala-tRNA^Pro^ with similar efficiency as bacterial Ala-tRNA^Pro^ (Fig. 1C). We hypothesize that differences in the acceptor stem sequences of these two tRNAs (Fig. 1A) may account for these differences in deacylation activity.

**Figure 2.**
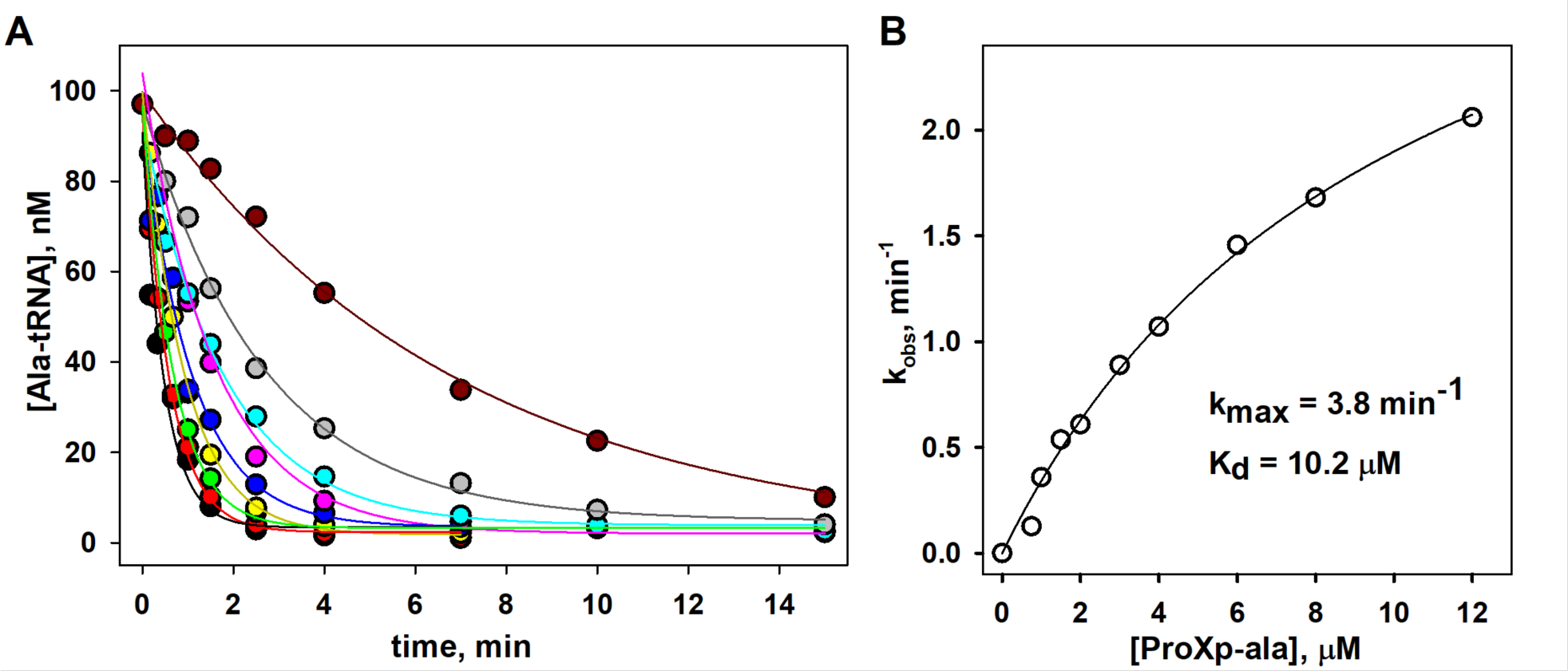
Deacylation of mischarged *Hs* cytosolic Ala-tRNA^Pro^ by *Hs* ProXp-ala under single-turnover conditions. (A) Time course of Ala-tRNA^Pro^ deacylation with varying concentration (0.75, 1.0, 1.5, 2.0, 3.0, 4.0, 6.0, 8.0 and 12 µM) of *Hs* ProXp-ala. Reactions were performed with 100 nM *Hs* Ala-tRNA^Pro^ at 30 °C. Deacylation curves were fit to a single exponential equation. (B) Hyperbolic fit of the observed rate constant vs *Hs* ProXp-ala concentration yielded a *k*_max_ of 3.8 min^-1^ and a K_d, app_ of 10.2 µM.

### Human ProXp-ala specificity is determined by tRNA acceptor stem elements

We previously showed that bacterial ProXp-ala cannot efficiently deacylate *Hs* cyto Ala-tRNA^Pro^ because of the differences in the identities of the bases at position 72 and 73 in this eukaryotic tRNA^Pro^ (Fig. 1C) (26). To test whether *Hs* ProXp-ala exhibits cross-species deacylation capability, assays with *Ec* Ala-tRNA^Pro^ were performed under single-turnover conditions. We found that *Hs* ProXp-ala only weakly deacylated *Ec* Ala-tRNA^Pro^ with a 66-fold decrease in *k*_obs_ (Fig. 3 and Table 1). This is consistent with the observation that *Hs* mito Ala-tRNA^Pro^ (which shares the C1:G72 and A73 acceptor stem elements of *Ec* tRNA^Pro^) is a poor substrate for *Hs* ProXp-ala (Fig. 1B). To identify the tRNA features that determine the specificity of *Hs* ProXp-ala, we prepared three *Hs* cyto tRNA^Pro^ acceptor-stem mutants (C73A, double mutant G1C:C72G, and triple mutant G1C:C72G/C73A) that mimic the acceptor stem of *Ec* tRNA^Pro^ (Fig. 1A). We found that the C73A and G1C:C72G tRNA^Pro^ mutants were hydrolyzed 13- and 16-fold slower than WT Ala-tRNA^Pro^, respectively, whereas the triple mutant G1C:C72G/C73A tRNA^Pro^ was deacylated at an ∼100-fold reduced rate, similar to the decrease observed with *Ec* WT tRNA^Pro^ (Fig. 3 and Table 1). These data indicate that the first base pair and discriminator base of tRNA^Pro^ are important elements for *Hs* ProXp-ala recognition and deacylation activity. To further validate the role of these bases, we prepared a series of *Ec* tRNA^Pro^ acceptor-stem variants containing *Hs* tRNA^Pro^ elements (Fig. 1A). We found that *Ec* C73 and G1:C72 tRNA^Pro^ mutants were significantly better substrate for the human enzyme than *Ec* WT tRNA^Pro^, while *Ec* G1:C72/A73 tRNA^Pro^ was deacylated slightly more efficiently than *Hs* WT tRNA^Pro^ (Fig. 3 and Table 1). Together, these results show that the deacylation activity of *Hs* ProXp-ala depends strongly on the identity of the nucleotides at the top of the acceptor stem of *Hs* cyto tRNA^Pro^.

**Table 1.** Rate constants (*k*_obs_) for deacylation of Ala-tRNA variants by *Hs* ProXp-ala.

| tRNA | Acceptor stem | $k_{\text{obs}}$ (min<sup>-1</sup>) | Fold change |
| --- | --- | --- | --- |
| *Hs* tRNA<sup>Pro</sup> WT | G<sup>1</sup>:C<sup>72</sup>/C<sup>73</sup> | 0.791 ± 0.12 | - |
| *Hs* tRNA<sup>Pro</sup> A73 | G<sup>1</sup>:C<sup>72</sup>/A<sup>73</sup> | 0.059 ± 0.007 | 13 |
| *Hs* tRNA<sup>Pro</sup> C1:G72 | C<sup>1</sup>:G<sup>72</sup>/C<sup>73</sup> | 0.049 ± 0.002 | 16 |
| *Hs* tRNA<sup>Pro</sup> C1:G72/A73 | C<sup>1</sup>:G<sup>72</sup>/A<sup>73</sup> | 0.007 ± 0.001 | 113 |
| *Hs* mito tRNA<sup>Pro</sup> | C<sup>1</sup>:G<sup>72</sup>/A<sup>73</sup> | 0.019 ± 0.005 | 42 |
| *Ec* tRNA<sup>Pro</sup> WT | C<sup>1</sup>:G<sup>72</sup>/A<sup>73</sup> | 0.012 ± 0.001 | 66 |
| *Ec* tRNA<sup>Pro</sup> C73 | C<sup>1</sup>:G<sup>72</sup>/C<sup>73</sup> | 0.071 ± 0.019 | 11 |
| *Ec* tRNA<sup>Pro</sup> G1:C72 | G<sup>1</sup>:C<sup>72</sup>/A<sup>73</sup> | 0.131 ± 0.009 | 6 |
| *Ec* tRNA<sup>Pro</sup> G1:C72/C73 | G<sup>1</sup>:C<sup>72</sup>/C<sup>73</sup> | 1.20 ± 0.16 | 0.66 |
| *Ec* tRNA<sup>Ser</sup> | G<sup>1</sup>:C<sup>72</sup>/G<sup>73</sup> | 0.018 ± 0.001 | 44 |
| *Ec* tRNA<sup>Ala</sup> | G<sup>1</sup>:C<sup>72</sup>/A<sup>73</sup> | 0.055 ± 0.002 | 14 |
| *Ec* tRNA<sup>Cys</sup> | G<sup>1</sup>:C<sup>72</sup>/U<sup>73</sup> | 0.098 ± 0.009 | 8 |

Deacylation assays were performed under single-turnover conditions at 30 °C using 1.5 μM *Hs* ProXp-ala and 100 nM Ala-tRNA. All fold changes are relative to WT *Hs* tRNA<sup>Pro</sup>. Results are the average of three independent trials with the standard deviation indicated.

**Figure 3.**
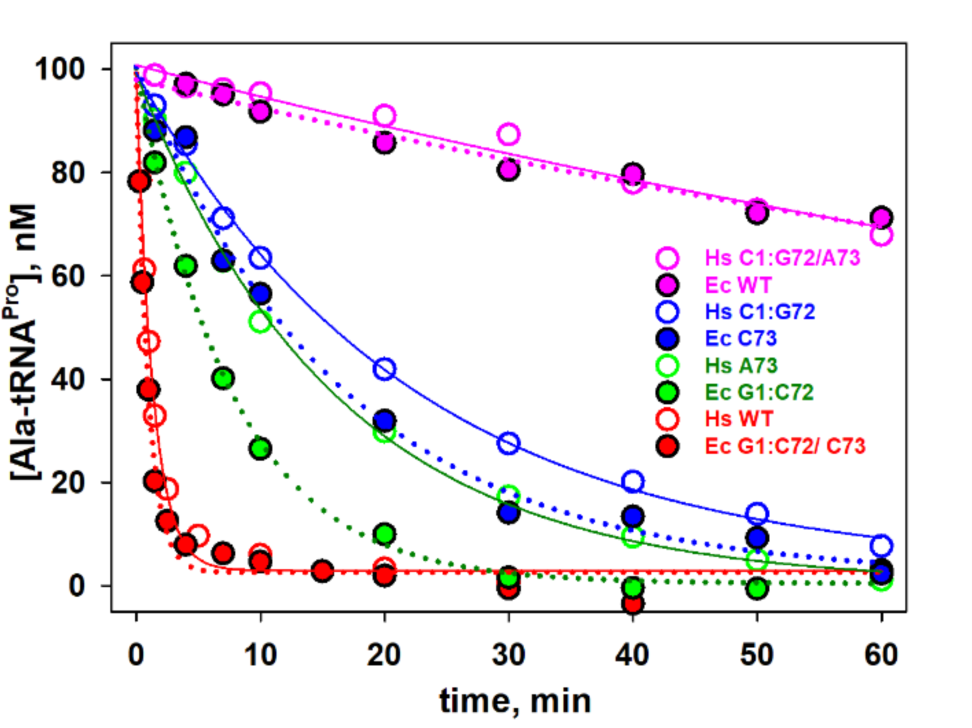
Deacylation of Ala-tRNA^Pro^ acceptor stem mutants by *Hs* ProXp-ala. *Hs* tRNA^Pro^ mutants (filled circles and solid fitting curves) and *Ec* tRNA^Pro^ variants (open circles and dashed fitting curves). The same color was chosen for tRNA^Pro^ mutants with identical nucleotides at positions 1:72 and 73: *Hs* WT and *Ec* G1:C72/C73 (red), *Hs* A73 and *Ec* G1:C72 (green), *Hs* C1:G72 and *Ec* C73 (blue), *Hs* C1:G72/A73 and *Ec* WT (magenta). Reactions were performed with 1.5 µM *Hs* ProXp-ala and 100 nM Ala-tRNA^Pro^ at 30 °C. Representative time courses of 3 independent experiments are shown.

To further study the effect of different substitutions at the 73 position, we charged or mischarged three *Ec* tRNAs with Ala: tRNA^Ala^, tRNA^Ser^ and tRNA^Cys^, which contain A73, G73 and U73, respectively (all have G1:C72). We found that the efficiency of Ala-tRNA^Ala^ deacylation is very similar to other G1:C73 tRNA^Pro^ variants with A73 (Table 1). Ala-tRNA^Cys^ (U73) was a slightly better substrate and Ala-tRNA^Ser^ (G73) displayed the lowest *k*_obs_ value of the three. These results emphasize a key role of the discriminator base for tRNA deacylation by *Hs* ProXp-ala.

### Aminoacyl substrate specificity of Hs ProXp-ala

We next investigated the aminoacyl specificity of *Hs* ProXp-ala using *Hs* tRNA^Pro^ charged with amino acids of different classes including 2-aminobutyric acid (Abu), Ser, Thr, Pro, and azetidine-2-carboxylic acid (Aze) (Fig. 4). The deacylation rate of tRNA^Pro^ charged with the larger non-polar Abu amino acid is 20-fold slower than Ala-tRNA^Pro^ (Table 2). Likewise, for tRNA^Pro^ charged with polar amino acids, the larger Thr-tRNA^Pro^ is a poorer substrate than Ser-tRNA^Pro^. Thus, for these amino acids, the deacylation efficiency correlates with size of the aminoacyl moiety, with a preference for the smaller substrate. In contrast, Aze, a 4-membered ring analog of Pro, with a molecular volume slightly larger than Ser, was deacylated 6-fold more efficiently than Ser and only 3.4-fold less efficiently than Ala. Aze-tRNA^Pro^ deacylation is also slightly greater (∼2-fold) than the rate of Pro-tRNA^Pro^ deacylation by the human enzyme, which surprisingly is only ∼5-fold lower than that of Ala-tRNA^Pro^ (Table 2).

**Table 2.** Rate constants (*k*_obs_) for deacylation of *Hs* aa-tRNA^Pro^ by *Hs* ProXp-ala.

| Aminoacyl-tRNA<sup>Pro</sup> | Amino Acid Volume (Å<sup>3</sup>) | $k_{\text{obs}}$ (min<sup>-1</sup>) | Fold change |
| --- | --- | --- | --- |
| Ala-tRNA<sup>Pro</sup> | 101 | $0.791 \pm 0.12$ | - |
| Abu-tRNA<sup>Pro</sup> | 122 | $0.039 \pm 0.007$ | 20 |
| Ser-tRNA<sup>Pro</sup> | 110 | $0.039 \pm 0.003$ | 20 |
| Thr-tRNA<sup>Pro</sup> | 129 | $0.015 \pm 0.002$ | 53 |
| Aze-tRNA<sup>Pro</sup> | 117 | $0.233 \pm 0.026$ | 3.4 |
| Pro-tRNA<sup>Pro</sup> | 132 | $0.158 \pm 0.012$ | 5.0 |

Deacylation assays were performed at 30 °C using 1.5 μM *Hs* ProXp-ala and 100 nM aa-tRNA<sup>Pro</sup>. Fold change is reported relative to *Hs* Ala-tRNA<sup>Pro</sup>. Results are the average of three independent trials with the standard deviation indicated.

**Figure 4.**
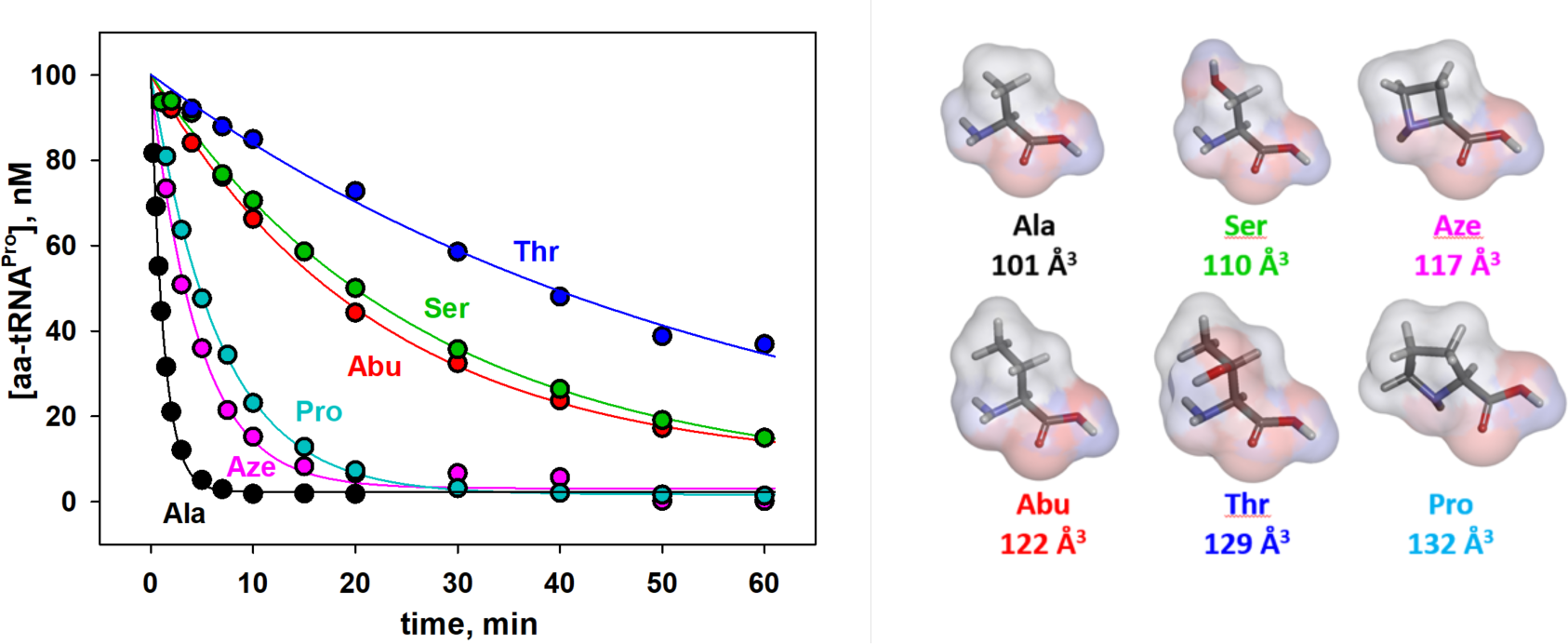
*Hs* ProXp-ala deacylation of *Hs* tRNA^Pro^ charged with various amino acids. Deacylation of Ala-tRNA^Pro^ (black), Aze-tRNA^Pro^ (magenta), Pro-tRNA^Pro^ (cyan), Abu-tRNA^Pro^ (red), Ser-tRNA^Pro^ (green), and Thr-tRNA^Pro^ (blue). Reactions were performed with 1.5 µM *Hs* ProXp-ala and 100 nM *Hs* aa-tRNA^Pro^ at 30 °C. Representative time courses of 3 independent experiments are shown. The structure and size of each amino acid, calculated as described in the Experimental Procedures, is shown on the right.

### The amino acid specificity of Hs ProXp-ala is tunable

The aminoacyl specificity of the *Ec* INS editing domain is modulated by conserved residues in the substrate binding pocket that include L266 and H369 (18,25). Whether the corresponding residues play a role in determining the overall specificity of *Hs* ProXp-ala is unknown. L266 and H369 of *Ec* INS align with M35 and H137 of *Hs* ProXp-ala, respectively (Fig. 5). We previously showed that L266 plays a role in the specificity of the INS towards Ser-tRNA; when this residue in *Ec* INS is mutated to Ala, Ala-tRNA^Pro^ is deacylated less efficiently, while the mutant enzyme hydrolyzes Ser-tRNA^Pro^ more robustly than WT INS (25). We now show that replacing M35 of *Hs* ProXp-ala with smaller Gly increases the rate of Ser-tRNA^Pro^ deacylation by 2.3-fold while decreasing the rate of Ala-tRNA^Pro^ deacylation by 4.7-fold (Fig. 6 and Table 3). The highly conserved His residue in the substrate binding pocket of *Ec* INS has previously been shown to function to prevent cognate substrate deacylation; mutation of this residue to Ala significantly increased Pro-tRNA^Pro^ deacylation (18). We now show that a homologous mutation (H137A) in *Hs* ProXp-ala also improves its activity towards Pro-tRNA^Pro^, which is now deacylated only 2-fold weaker than Ala-tRNA^Pro^ by the same mutant (Fig. 7A and Table 3). Similarly, a *Cc* H130A ProXp-ala variant showed strong deacylation of Pro-tRNA^Pro^ (Fig. 7B). Interestingly, the Ala-tRNA^Pro^ deacylation activities of *Cc* H130A and *Hs* H137A ProXp-ala were not significantly different than those of the WT enzymes.

**Table 3.** Rate constants (*k*_obs_) for deacylation of *Hs* aa-tRNA^Pro^ by WT and mutant *Hs* ProXp-ala.

|  | **Ala-tRNA<sup>Pro</sup>** | **Ser-tRNA<sup>Pro</sup>** |  | **Pro-tRNA<sup>Pro</sup>** |  |
| --- | --- | --- | --- | --- | --- |
| | $k_{\text{obs}}$ (min<sup>-1</sup>) | $k_{\text{obs}}$ (min<sup>-1</sup>) | Fold change<br>(Ala/Ser) | $k_{\text{obs}}$ (min<sup>-1</sup>) | Fold change<br>(Ala/Pro) |
| **WT** | 0.791 ± 0.12 | 0.039 ± 0.003 | 20 | 0.158 ± 0.012 | 5.0 |
| **M35G** | 0.170 ± 0.008 | 0.089 ± 0.011 | 1.9 | 0.029 | 5.9 |
| **H137A** | 0.804 ± 0.019 | 0.045 ± 0.004 | 18 | 0.381 ± 0.056 | 2.1 |

Deacylation assays were performed at 30 °C using 1.5 μM *Hs* ProXp-ala and 100 nM *Hs* aa-tRNA<sup>Pro</sup>. With the exception of Pro-tRNA<sup>Pro</sup> deacylation by *Hs* ProXp-ala M35G (1 trial), all results are the average of three independent trials with the standard deviation indicated.

**Figure 5.**
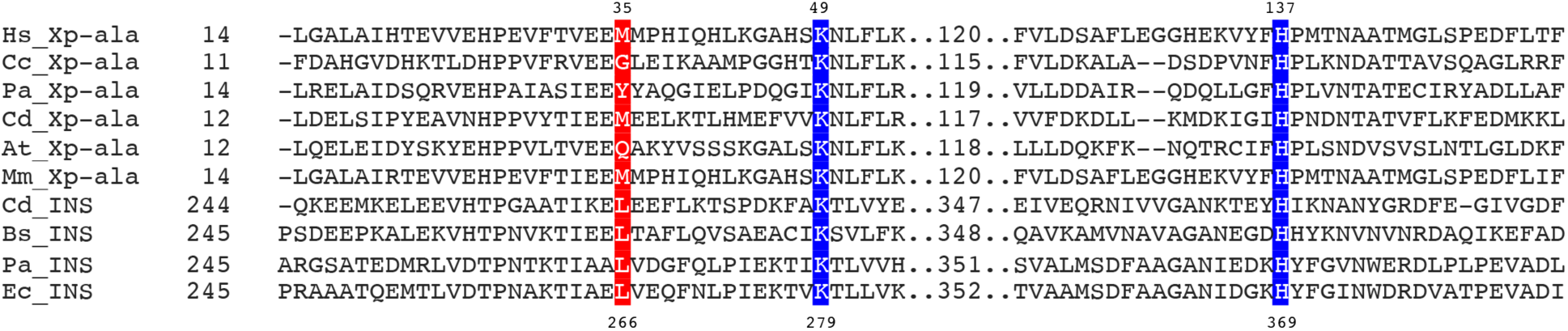
Multiple sequence alignment of ProRS INS and ProXp-ala. Sequences of *E. coli* (Ec), *Clostridioides difficile* (Cd), *Bacillus subtilis* (Bs), and *Pseudomonas aeruginosa* (Pa) INS domains were aligned with *Homo sapiens* (Hs), *Mus Musculus* (Mm), *Arabidopsis thaliana* (At), *C. crescentus* (Cc), *P. aeruginosa* (Pa), and *C. difficile* (Cd) ProXp-ala sequences using Clustal Omega (47). Corrections to the resulting alignment were made based on the structural alignment of *Cc* ProXp-ala (5VXB) and *Pa* ProRS (5UCM). Strictly and partially conserved residues within the ProXp-ala and INS families investigated here are highlighted in blue and red, respectively. Numbered residues correspond to the sequences of *Ec* INS and *Hs* ProXp-ala, respectively.

**Figure 6.**
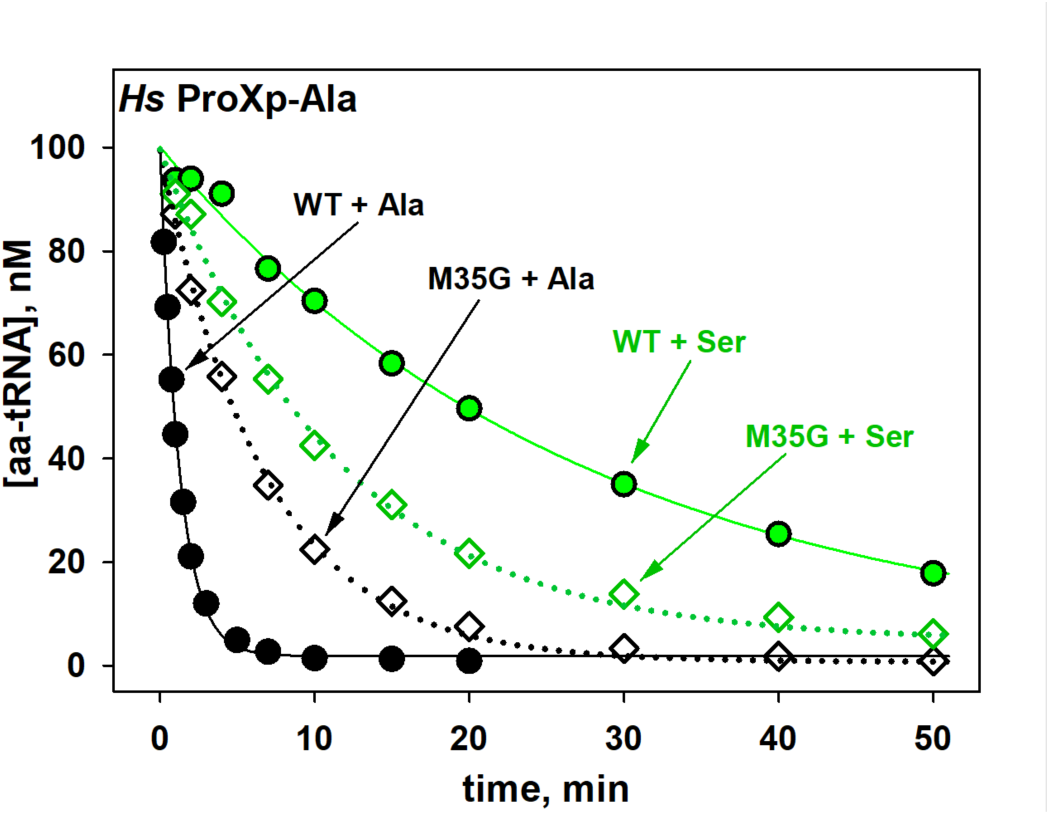
Deacylation of *Hs* Ser- and Ala-tRNA^Pro^ by WT and mutant ProXp-ala. Deacylation of tRNA^Pro^ mischarged with Ala (black) and Ser (green) by WT (circles) and M35G (diamonds) *Hs* ProXp-ala. Reactions were performed with 1.5 µM *Hs* ProXp-ala and 100 nM *Hs* aminoacyl-tRNA^Pro^ at 30 °C. Representative time courses of 3 independent experiments are shown.

**Figure 7.**
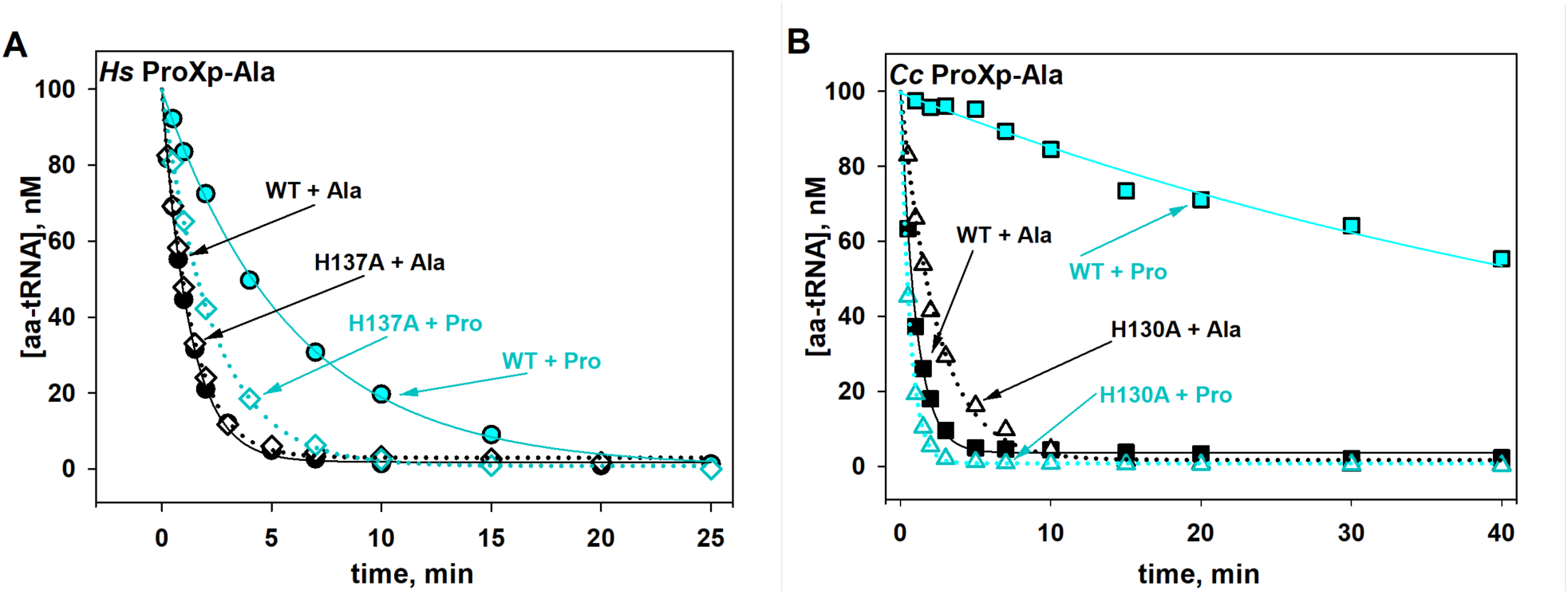
Deacylation of Ala- and Pro-tRNA^Pro^ by WT and mutant ProXp-ala. (A) Deacylation of *Hs* tRNA^Pro^ charged with Ala (black) and Pro (cyan) by WT (circles) and H137A (diamonds) *Hs* ProXp-ala. (B) Deacylation of *Ec* tRNA^Pro^ charged with Ala (black) and Pro (cyan) by WT (squares) and H130A (triangles) *Cc* ProXp-ala. Reactions were performed with 1.5 µM *Hs* ProXp-ala and 100 nM *Hs* aminoacyl-tRNA^Pro^ at 30 °C or with 0.75 µM *Cc* ProXp-ala and 100 nM *Ec* aminoacyl-tRNA^Pro^ at 18 °C. Representative time courses of 3 independent experiments are shown.

### Phylogeny and distribution of the ProXp-ala family

Previous bioinformatic analyses showed that bacterial ProXp-ala is primarily encoded in organisms belonging to the Alphaproteobacteria, Lactobacillales, Clostridia, and Negativicutes classes (20,22), whereas eukaryotic ProXp-ala have only been reported for *Hs, Mus musculus, Arabidopsis thaliana, Trypanosoma brucei* and *Plasmodium falciparum* (20). To update our understanding of the overall phylogenetic distribution of ProXp-ala in Eukarya, we searched all available 540 eukaryotic sequenced genomes and identified ProXp-ala genes in approximately 54% of the organisms analyzed (Table S1, Fig. 8). ProXp-ala is widely distributed in mammals, birds, fish, reptiles and amphibians, while it is absent in other analyzed Animalia groups such as insects, arthropods, and worms. ProXp-ala is also encoded by almost all sequenced plants. The predicted plant ProXp-ala gene sequence is almost twice the size (∼320 amino acids) of that found in bacteria and the Animalia kingdom. The novel C-terminal extension of plant ProXp-ala shares no primary sequence homology with any known protein and its function is unknown. In contrast to plants and animals, ProXp-ala is fused to the N-terminus of ProRS in a group of animal parasites belonging to the Stramenopila, Aveolates and Rhizaria (SAR) supergroup, which includes *P. falciparum* and *Toxoplasma gondii*. This ProXp-ala-ProRS fusion is also observed in *Trypanosoma brucei* ProRS and some other species from Kinetoplasida (e.g., Leishmania). We also identified putative ProXp-ala genes in several archaeal species including members of the Asgard archaea superphylum, *Candidatus* Prometheoarchaeum syntrophicum MK-D1 and *Lokiarchaeum sp*. GC14_75. Both organisms have been associated with eukaryogenesis and may be provide a link to the evolution of ProXp-ala (33,34).

**Figure 8.**
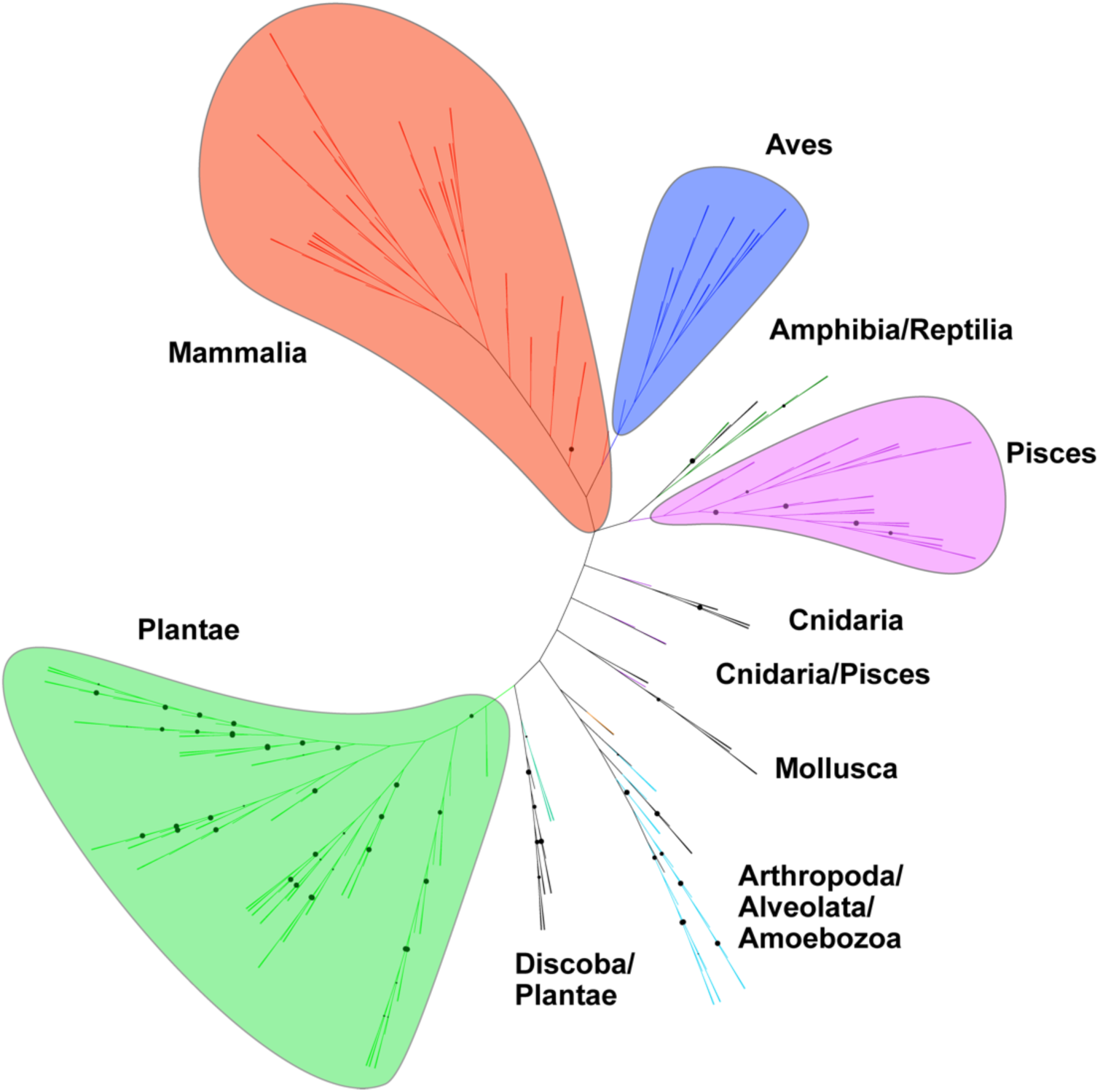
Unrooted phylogenetic tree of eukaryotic ProXp-ala. ProXp-ala genes are primarily encoded in organisms belonging to the Animalia (Mammalia, Aves, Amphibia, Reptilia, Pisces, Cnidaria, Mollusca, and Arthropoda) and Plantae kingdoms as well as in the Alveolata, Discoba, and Amoebozoa clades. Branches with bootstrap values higher than 90 are indicated by black circles.

A phylogenetic tree of the ProXp-ala family revealed the formation of distinct clades corresponding to eukaryotic and bacterial ProXp-ala with only two instances of possible horizontal gene transfer events in the eukaryotes *Entamoeba dispar* and *Ostreococcus tauri* (Fig. 9). The phylogenic relationship of the ProXp-ala family underlines their divergence during evolution, which is reflected in distinct substrate features.

**Figure 9.**
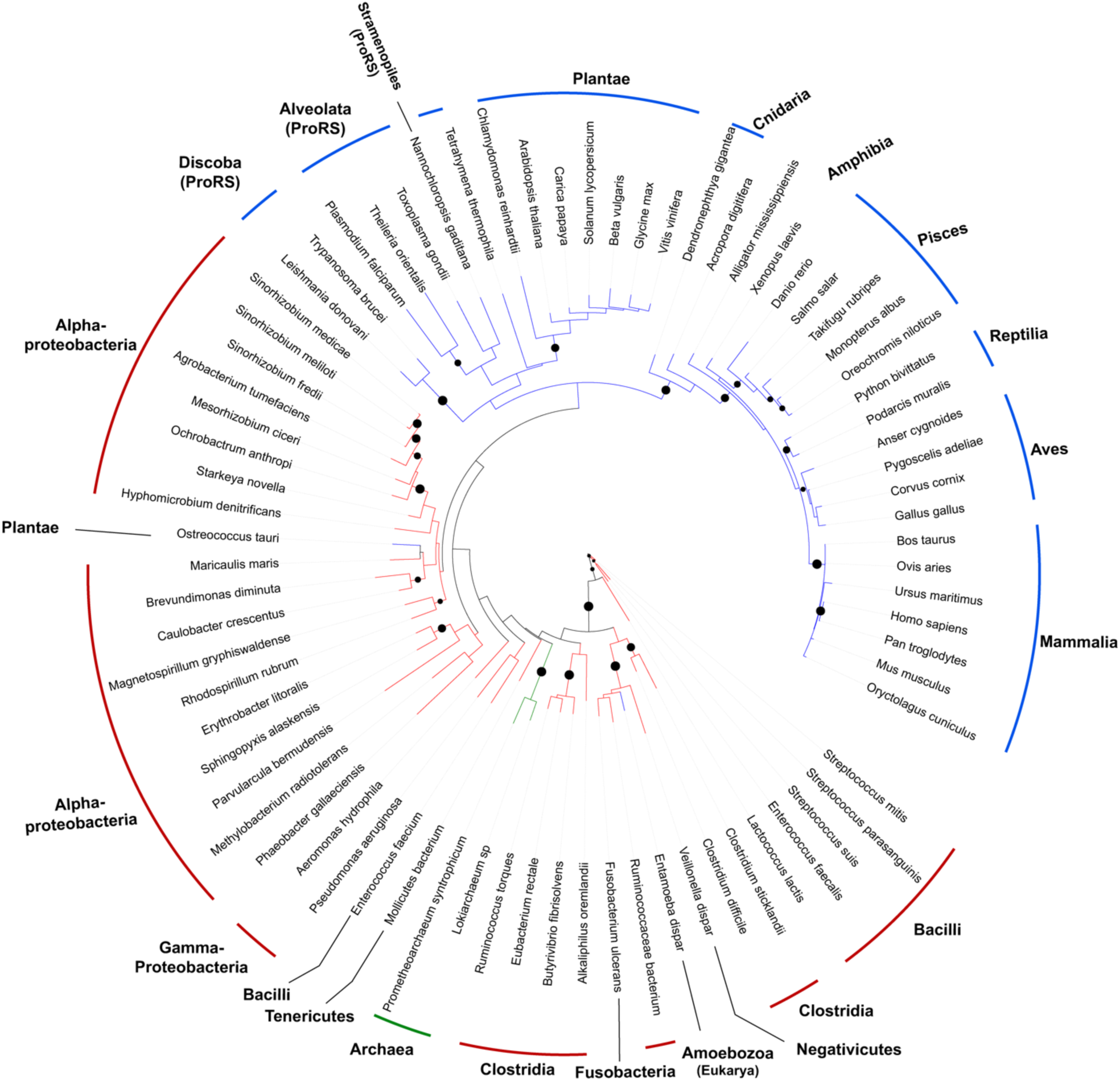
Phylogeny of ProXp-ala. The branches corresponding to bacterial, eukaryotic, and archaeal ProXp-ala are colored in red, blue, and green, respectively. The phylum, clade, or kingdom to which each organism belongs are indicated. ProXp-ala is fused to ProRS in organisms from the Discoba, Alveolata, and Stramenopiles clades. Branches with bootstrap values higher than 70 are indicated by black dots.

## Discussion

In contrast to a previous report, which concluded that *Hs* cyto ProXp-ala lacks tRNA specificity (31), we found a strong dependence on elements at the top of the acceptor stem for efficient deacylation by the *Hs* enzyme (Table 1). In the earlier study, the reported activity was weak even at an elevated enzyme concentration of 5 µM (31). It is possible that the relatively weak deacylation activity was due to the use of a tRNA^Pro^ mutant with three substitutions (C3G, G70U, and C73A) relative to WT tRNA^Pro^. These mutations were introduced to facilitate the preparation of Ala-tRNA^Pro^ using AlaRS (31). While we did not test the third base pair in our work directly, we found that C73 is a critical recognition element for *Hs* ProXp-ala and G3:U70-containing *Ec* Ala-tRNA^Ala^ is a relatively poor substrate despite the fact that it contains G1:C72 (Table 1). In addition, differences in the N-terminus of the recombinant *Hs* ProXp-ala used previously relative to the genome-encoded sequence available in UniProt (35) may have impacted enzymatic function (Fig. S2) (31).

The changes in the sequence of the acceptor stem of tRNA^Pro^ during evolution led to the development of distinct modes of tRNA recognition by ProRSs across the three domains of life (28,30,36). Here we found that these same changes also exerted pressure during the evolution of the *trans*-editing enzyme ProXp-ala from Bacteria to Eukarya, enabling its adaptation to cytosolic tRNA^Pro^. However, *Hs* ProRS and ProXp-ala altered their substrate selection mechanisms in response to the tRNA^Pro^ acceptor stem changes differently. *Hs* ProRS relaxed its requirement for specific acceptor stem nucleotides and instead gained a stronger affinity for the anticodon, as well as a specific D-stem loop requirement (30). In contrast, *Hs* ProXp-ala specificity relies on the G1:C72/C73 nucleotides of tRNA^Pro^ (Fig. 3, Table 1), which are the same acceptor stem positions recognized by bacterial ProXp-ala, albeit with different base identities (*i*.*e*. C1:G72/A73) (26). Consequently, *Hs* ProXp-ala only weakly deacylated bacterial and mitochondrial Ala-tRNA^Pro^ (Fig. 1B), and its strong tRNA specificity likely prevents efficient deacylation of Ala-tRNA^Ala^ (Table 1). These results, and the fact that C73, which is rarely found at this position and is a universal feature of all eukaryotic cyto tRNA^Pro^, indicate that ProXp-ala is likely to act exclusively in the cytosol of higher eukaryotes. Studies to determine the cellular localization of *Hs* ProXp-ala are required to confirm this hypothesis. Thus, whereas acceptor stem elements of bacterial tRNA^Pro^ play a dual role in aminoacylation and editing, the acceptor stem elements that only weakly mark cytosolic tRNA^Pro^ for aminoacylation with Pro by *Hs* ProRS, play a more important role in the proofreading of aminoacyl-tRNA^Pro^ by *Hs* ProXp-ala.

Biochemical characterization also showed that *Hs* ProXp-ala displays an aminoacyl specificity profile that does not follow a simple relationship between the rate of hydrolysis and the size (molecular volume) of the amino acid. For example, *Hs* ProXp-ala deacylated Abu-tRNA^Pro^ with similar efficiency as Ser-tRNA^Pro^, despite Ser being significantly smaller than Abu. In contrast, tRNA^Pro^ charged with Aze (which resembles the volume of Ser) was deacylated several fold better than Ser-tRNA^Pro^ and with similar efficiency as Pro-tRNA^Pro^ (Fig. 4, Table 2). Thus, in comparison to the bacterial system (24), the *Hs* ProXp-ala active site better accommodates the pyrrolidine and azetidine side chains of Pro and Aze and shows greater discrimination against polar OH-containing amino acids. Consequently, *Cc* and *Hs* ProXp-ala appear to have evolved not only different tRNA acceptor stem recognition specificities but also different preferences in amino acid selection.

Our study uncovered key residues that define the amino acid specificity of the ProXp-ala family. In *Hs* ProXp-ala, mutations at M35 and H137 modulated amino acid recognition, similar to the corresponding Leu and His residues in the substrate binding pocket of the *Ec* INS domain (25). Substitution of M35 with G enlarges the *Hs* enzyme’s binding pocket and allowed better accommodation of Ser-tRNA^Pro^. The corresponding position in *Cc* ProXp-ala corresponds to G32, and WT *Cc* ProXp-ala has a binding pocket that enables it to efficiently accommodate and robustly hydrolyze Ser-tRNA^Pro^ (24).

Our results also confirmed that a highly conserved His residue in the aminoacyl-binding pocket of INS/ProXp-ala family members functions as a universal gatekeeper to prevent or reduce cognate Pro-tRNA^Pro^ deacylation. In the case of INS, the H369A mutation led to a switch of specificity from Ala-to Pro-tRNA^Pro^ (18), whereas the analogous mutation in *Cc* and *Hs* ProXp-ala resulted in dual activity for Ala- and Pro-tRNA^Pro^. Notably, H137 of *Hs* ProXp-ala is less effective at preventing Pro-tRNA^Pro^ deacylation relative to the bacterial system (Fig. 7). The rather efficient deacylation of cognate Pro-tRNA^Pro^ observed for the *Hs trans*-editing enzyme *in vitro* is surprising and suggests that other factors in the cell likely prevent this undesired reaction. Based on the ability of bacterial elongation factor Tu (EF-Tu) to protect correctly charged tRNAs from free-standing editing domains, including bacterial ProXp-ala (22,37), we propose that *Hs* EF-1A may outcompete *Hs* ProXp-ala for binding to Pro-tRNA^Pro^, allowing only deacylation of the mischarged Ala-tRNA^Pro^ species in cells. If editing occurred in *cis* (i.e., by a ProRS editing domain prior to substrate dissociation) rather than in *trans*, this competition would be less likely to be successful and may explain why eukaryotic ProRSs rely on a free-standing *trans*-editing enzyme for proofreading. Whether Aze-tRNA^Pro^ is protected by elongation factors is unknown. Aze is a proline analog synthesized in many plants; it is also misactivated by *Hs* ProRS (15). High levels of Aze administered to zebrafish embryos lead to Aze-tRNA^Pro^ accumulation and cell toxicity (38). Thus, it is possible that the dual activity towards Ala and Aze, albeit weaker for the latter, displayed by *Hs* ProXp-ala serves to prevent mistranslation of Pro codons with either amino acid when present at low levels.

The results reported here underscore the complex evolutionary relationship between class II aminoacyl-tRNA synthetases and editing domains. The tRNA^Pro^-specific INS/ProXp-ala family of editing domains likely arose from the biological pressure to prevent mistranslation of Pro codons due to the critical nature of this amino acid for protein structure and function in all Domains of life. The ProRS INS domain, found only in bacteria (14,39), and the ProXp-ala family, found primarily in bacteria and eukarya (Fig. 9), have the same specificity toward editing of Ala-tRNA^Pro^; yet, ProXp-ala is usually encoded as a free-standing domain (with a few exceptions, see Fig. 9) and INS is always found appended to ProRS (14,20,22,39). The latter observation may be due to the lack of strong tRNA acceptor stem specificity of this domain (26), which would result in promiscuous *trans*-editing of Ala-tRNA^Ala^. Each organism has likely evolved to use the type of Ala-tRNA^Pro^ editing domain (INS vs. ProXp-ala) and domain structure (free-standing or synthetase-appended) to ensure optimal levels of fidelity required for cell survival.

## Experimental procedures

### Materials

All amino acids, nucleotides, enzymes and other materials were purchased from MilliporeSigma unless otherwise noted.

### Protein preparation

All WT and mutant proteins used in this study were prepared as previously described: *Hs* ProXp-Ala (38), *Cc* ProXp-Ala (22), *Hs* ProRS (40), *Ec* ProRS K279A (25), *Ec* tRNA nucleotidyltransferase (5). Briefly, BL-21-CodonPlus(DE3)-RIL cells (Agilent) were used for protein expression. Overexpression was carried out upon induction with 0.1 mM isopropyl β-D-1-thiogalactopyranoside (Gold Biotechnology) for 18-20 hours at 18 °C. His6-tagged proteins were purified on His-Select Nickel Affinity Gel using a 5-250 mM imidazole gradient. Purified proteins were buffer exchanged into the storage buffer (50 mM sodium phosphate pH 7.5, 150 mM NaCl, 1 mM DTT), mixed with 40% glycerol and stored at -20 °C. Enzyme concentrations were determined using a Bio-Rad Protein Assay Kit (Bio-Rad). Mutations in *Hs* ProXp-ala and *Cc* ProXp-Ala were introduced using either QuikChange Site-Directed Mutagenesis (Agilent) or Site-directed, Ligase-Independent Mutagenesis (SLIM) (41). The successful mutagenesis was confirmed by DNA sequencing (The Genomics Shared Resource at The Ohio State University Comprehensive Cancer Center).

### Preparation of tRNAs and aminoacyl-tRNA substrates

All tRNA variants used in these studies were prepared by *in vitro* transcription as previously described (14). Briefly, for each tRNA, a linear DNA template for T7 RNA polymerase was obtained by BstN1 (NEB) digestion of a plasmid carrying the corresponding gene. Transcribed tRNA was purified using denaturing 10% polyacrylamide gel electrophoresis. To ensure high yields of tRNA variants with cytidine at position 1, a hammerhead ribozyme was inserted at 5′ of the tRNA gene (42). A plasmid encoding *Hs* tRNA^Pro^ C1:72G variant was ordered from Genewiz; all other acceptor stem tRNA^Pro^ mutants were made by SLIM (41) and confirmed by DNA sequencing (The Genomics Shared Resource at The Ohio State University Comprehensive Cancer Center). All tRNAs used in deacylation assays were labeled with ^32^P at the 3′-A76 using tRNA nucleotidyltransferase (43). Preparation of aminoacylated *Hs* tRNA^Pro^ was carried out in 50 mM HEPES, pH 7.5, 20 mM KCl, 4 mM ATP, 10 mM MgCl_2_, 0.1 mg/mL BSA, and 1 mM DTT by incubating 10 µM *Hs* ProRS, 10 µM tRNA^Pro^, and 0.03 mg/mL pyrophosphatase (Roche) with the following amino acids: 900 mM Ala, 30 mM Pro, or 300 mM Aze for 5 min at 37 °C. *Ec* Pro- and Ala-tRNA^Pro^ were prepared under similar conditions with editing deficient *Ec* K279A ProRS (25). Aminoacyl-tRNAs were phenol chloroform-extracted followed by ethanol precipitation. Acceptor-stem tRNA^Pro^ variants as well as *Ec* tRNA^Ser^, *Ec* tRNA^Ala^, and *Ec* tRNA^Cys^ were charged with Ala using dinitro-flexizyme (dFx) and Ala-3,5-dinitrobenzyl ester (Ala-DBE) as described (44). Briefly, ^32^P-labeled tRNA and dFx were refolded by heating at 90 °C for 1 min followed by addition of MgCl_2_ and cooling to room temperature for 3 min. The aminoacylation reaction was carried in 100 mM HEPES-KOH (pH 7.5), 20 mM MgCl_2_, 25 µM tRNA, 25 µM dFx, 5 mM Ala-DBE, Ser-DBE or Abu-DBE for 20 h on ice. Thr-tRNA^Pro^ was prepared using enhanced-flexizyme (eFx) and Thr-4-chlorobenzyl thioester (44). Substrates for deacylation assays were ethanol precipitated and dissolved in 3 mM sodium acetate, pH 5.2, and stored at -80 °C prior to use.

### Deacylation assays

Single-turnover aminoacyl-tRNA^Pro^ deacylation reactions were performed as previously described (24). *Hs* ProXp-Ala (WT and mutants) reactions typically contained 0.1 μM ^32^P-labeled aminoacyl-tRNA^Pro^ and 1.5 μM ProXp-ala in deacylation buffer A (50 mM HEPES pH 7.5, 20 mM KCl, 5 mM MgCl_2_, 0.1 mg/ml BSA, 1 mM DTT) and were performed at 30 °C. *Cc* ProXp-Ala reactions contained 0.1 μM ^32^P-labeled aminoacyl-tRNA^Pro^ and 0.75 μM ProXp-ala in deacylation buffer B (150 mM potassium phosphate pH 7.0, 5 mM MgCl_2_, 0.1 mg/mL BSA) and were performed at 18 °C. Reactions were initiated by mixing equal volumes of aminoacyl-tRNA^Pro^ and enzyme. At the chosen time points, 2-µL aliquots were quenched into a solution containing 20 units S1 nuclease, 1 mM zinc acetate, and 200 mM NaOAc mM sodium acetate, pH 5.0. Product formation was monitored by separating aminoacyl-A76 from A76 on PEI-cellulose plates (EMD Millipore) using a mobile phase of 0.05% ammonium chloride/5% acetic acid. Radioactive products were detected by autoradiography using a Typhoon FLA 9500 (GE Healthcare) and quantified using ImageQuant TL 8 software (GE Healthcare). Comparative analysis of numerous tRNA and enzyme variants was performed using single-turnover rate conditions with the concentration of *Hs* ProXp-ala several fold smaller than the apparent K_d_. Under these conditions, the observed rate constant (*k*_obs_) reflects *k*_cat_/*K*_m_ and allows comparison of the enzyme efficiency for various substrates. For K_d_ determination, deacylation reactions were performed with varying *Hs* ProXp-Ala concentrations from 0.75 µM to 12 µM. The K_d_ and *k*_max_ were obtained by fitting the *k*_obs_ vs [*Hs* ProXp-Ala] plot with the Michaelis–Menten equation: *k*_max_ = *k*_obs_ [*Hs* ProXp-Ala]/(K_d_ + [*Hs* ProXp-Ala]). Observed rate constants were obtained by fitting the time course for aminoacyl-tRNA deacylation with a single-exponential equation using SigmaPlot (Systat Software). Each reported rate constant is an average of three independent assays. The graphs show representative time courses. All data points were corrected for non-enzymatic buffer hydrolysis.

### Amino Acid Volume Calculations

The structure of each free amino acid (Ala, Abu, Aze, Pro, Ser, and Thr) was obtained from the Protein Data Bank. Molecular volumes for each amino acid were calculated by loading each .pdb file into the Volume Assessor module of the 3V webserver (http://3vee.molmovdb.org/volumeCalc.php) using a 0.1 Å probe radius and high grid resolution (0.5 Å voxel size) (45).

### Phylogenetic analysis of ProXp-ala

A total of 542 eukaryotic sequenced genomes (retrieved from the KEGG database, www.genome.jp, and the Joint Genome Institute Genomes Online Database (46)) were individually searched for ProXp-ala genes using the National Center for Biotechnology Information BLASTp service (Table S1). ProXp-ala reference sequences from *Hs, Cc, Clostridium sticklandii, Agrobacterium tumefaciens*, and *Arabidopsis thaliana* were used as queries. A global search of ProXp-ala genes in Archaea was also carried out. The resulting putative genes were designated as ProXp-ala only if they had the signature residues of ProXp-ala (*i*.*e*. K45, H130, GXXXP/A, numbering based on *C. crescentus* ProXp-ala). The eukaryotic ProXp-ala phylogenetic tree was built via multiple sequence alignment using Clustal Omega (47) followed by a Maximum Likelihood analysis with 100 bootstraps using MEGA X with default settings (48). The ProXp-ala family tree was built using the NGPhylogeny.fr platform (49). A workflow with Clustal Omega sequence alignment followed by tree inference using PhyML with 100 bootstraps was used. The bacterial, eukaryotic, and archaeal ProXp-ala sequences from the organisms listed in Table S1 were used for the tree. The iTOL online tool was used to display both trees (50).

## Acknowledgments

We thank Irina Shulgina for protein and tRNA preparation, Dr. Marie Sissler (Université de Strasbourg) for providing the plasmid encoding the human mitochondrial tRNA^Pro^, Dr. Ana Crnkovic for critical reading of the manuscript, and Drs. Mom Das and Sandeep Kumar for enlightening discussions and suggestions.

## Funding and additional information

This work was supported by NIH grant RO1 GM113656 to K.M-F and by KAKENHI (JP20H05618 to H.S. and JP20H02866 to Y.G.) from the Japan Society for the Promotion of Science.

## Abbreviations

The abbreviations used are:

*Cc*: *Caulobacter crescentus*
*Ec*: *Escherichia coli*
*Hs*: *Homo sapiens*
ARS: aminoacyl-tRNA synthetase
ProRS: prolyl-tRNA synthetase
AlaRS: alanyl-tRNA synthetase
EF-Tu: elongation factor Tu,
tRNA: transfer RNA
aa-tRNA: aminoacyl-tRNA
WT: wild-type
Abu: 2-aminobutyric acid
Aze: azetidine-2-carboxylic acid
cyto: cytosolic
INS: insertion domain
mito: mitochondrial
Abu: 2-aminobutyric acid

